# Thermodynamics of DNA Hybridization from Atomistic Simulations

**DOI:** 10.1101/2020.08.05.238485

**Authors:** Gül H. Zerze, Frank H. Stillinger, Pablo G. Debenedetti

**Affiliations:** Department of Chemical and Biological Engineering, Princeton University, Princeton, New Jersey 08544, USA; Department of Chemistry, Princeton University, Princeton, New Jersey 08544, USA

## Abstract

Studying the DNA hybridization equilibrium via brute force molecular dynamics (MD) or commonly used advanced sampling approches is notoriously difficult at atomistic lengthscale. However, besides providing a more realistic modeling of this microscopic phenomenon, atomistic resolution is a necessity for some fundamental research questions, such as the ones related to DNA’s chirality. Here, we describe an order parameter-based advanced sampling technique to calculate the free energy surface of hybridization and estimate melting temperature of DNA oligomers at atomistic resolution, using a native topology-based order parameter. We show that the melting temperatures estimated from our atomistic simulations follow an order consistent with the predictions from melting experiments and those from the nearest neighbor model, for a range of DNA sequences of different GC content. Moreover, free energy surfaces and melting temperatures are calculated to be identical for D- and L-enantiomers of Drew-Dickerson dodecamer.

Graphical TOC Entry

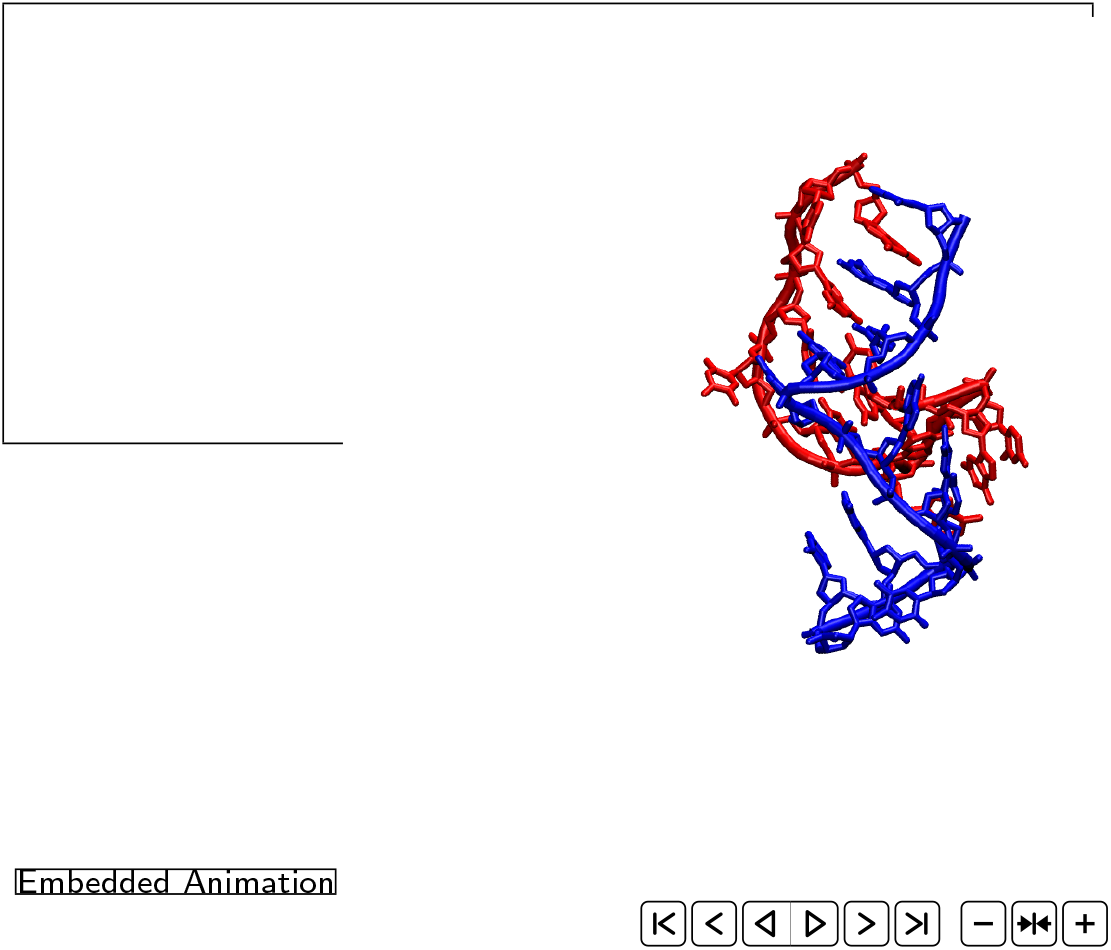

## Introduction

A fundamental biophysical process is DNA hybridization which involves noncovalent attachment of two strands of DNA chains to one another, resulting in a double-stranded helix (duplex). Double-stranded helix may unwind back to the coil-like single-stranded configurations of strands via, for example, thermal melting, as repeatedly happens during a polymerase chain reaction. Like many other biological phenomena observed in nature, DNA hybridization is a phase transition which is notoriously rare and slow to observe especially at the atomistic scale.

Accordingly, computer simulations of DNA duplexes employ modeling approaches at different length scales for different purposes. State-of-the-art simulation studies of DNA hybridization thermodynamics use coarse-grained models of DNA. Prior to the simulations, DNA hybridization thermodynamics have been studied substantially via experiments, especially thermal melting experiments, which have formed foundations of elegant theories of nucleic acid melting, such as the nearest-neighbor or next-nearest-neighbor models.^1–7^ These experimental studies have also helped simulation studies to develop coarse-grained potentials of nucleic acids for studying the hybridization thermodynamics.^8–13^ Coarse-grained models of nucleic acids typically incorporate base-stacking, base-pairing, and some backbone dihedral interactions. However, given the slow dynamics of DNA hybridization, resolution of even the most detailed coarse-grained models are limited to several beads per nucleotide. Atomistic models of nucleic acids, on the other hand, are more widely used to study fine structural details, configurational variation, and flexibility of double stranded helices as well as transitions between different forms of duplexes (i.e., structural polymorphism in double-stranded DNA).^14–19^

Atomistic models have also been used to study the formation of single base-pairs in double stranded DNA^20^ and base-stacking free energy^21^ although studying a full-length chain was still out of the reach at that time. More recently, Piana has presented the first atomistic molecular dynamics (MD) study where he studied helix-coil transition of short DNA sequences (hexamers)^22^ using an advanced-sampling MD, specifically, bias-exchange metadynamics technique using pairwise coordination between atoms involving in Watson-Crick (WC) pairs and phosphate backbone dihedral angles as the order parameters to advance the sampling. However, the expected stability of DNA hybrid has not been successfully reproduced. Here, we propose a novel advanced-sampling approach in order to study the DNA hybridization thermodynamics using atomistic models. In this combined advanced sampling technique, we employ a parallel tempering scheme in conjunction with metadynamics using an order parameter inspired by the protein folding and binding studies. We calculated the free energy surface and melting temperature of Drew-Dickerson Dodacamer (DDD) with our technique and found that our melting temperature estimates agree well with the predictions from the melting experiments and those from the nearest neighbor model. We further tested our proposed technique simulating decamers of various GC content and found that the melting temperature estimates from our technique match with the experimental estimates within ±15°C. We note that the predictive capability of our simulations is limited by the predictive capability of underlying potential energy function (CHARMM36 nucleic acids). Although this force field is commonly used in atomistic simulations, it is not specifically optimized for predicting the hybridization free energy. We emphasize that such a task, i.e. optimizing the force field for better prediction of the free energy, would not be possible without a proper sampling technique, as the lack of equilibrium sampling of hybridization hinders pinpointing the sources of potential problems in the model. Therefore, our proposed sampling technique is also a useful asset for future refinements of nucleic acids force fields, as well.

## Methods

### DNA systems and modeling

We study the hybridization of a dodecamer and seven decamers. We, first, show the validity of our approach using Drew-Dickerson dodecamer (DDD), also known as EcoRI dode-camer. ^23–25^ To further validate our method, we test a number decamers of systematically varied GC content, for which extensive experimental data is available ^26^ (sequences from 5’ to 3’ are: ATCAATCATA, TTGTAGTCAT, CCAACTTCTT, ATCGTCTGGA, CGATCTGCGA, GATGCGCTCG, GGGACCGCCT). Finally, we study a DDD variant, fully composed of L-nucleotides, i.e. complete mirror image structure of DDD to measure the impact of the total chiral inversion.

We setup two independent simulations of DDD, where the only difference is in the initial condition. All replicas of one of the simulations are initiated from a completely unhybridized (equilibrated) structure where two strands are far apart from each other whereas all replicas are initiated from the (equilibrated) crystal structure (PDB ID: 1BNA) in the other simulation. Initial coordinates for decamers are generated using the CHARMM program and internal coordinates (IC) table supplied in CHARMM36 force field (for CHARMM program).^27–29^ The chirally-inverted DDD is generated modifying the IC table of CHARMM36 force field for CHARMM program (modified IC parameters are available upon request).

Each DNA is solvated in a truncated octahedron box with 5.7 nm and 6 nm spaced faces, respectively, for decamers and DDD. Number of water molecules are adjusted to achieve approximately 1000 kg/m^3^ density. Simulation box is large enough not to allow chain A (or chain B) to interact with itself from any periodic image of the simulation box within the limits of nonbonded interaction cutoff distance.

DDD DNA is studied at 100 mM NaCl concentration, whereas decamers are studied at 69 mM NaCl concentration as given in the experimental conditions. ^26^ Salt concentration is adjusted by adding related number of Na^+^ and Cl^*−*^ ions to the solution after enough number of counterions are added to provide the electroneutrality of system. All DNA systems are modeled using CHARMM36 nucleic acid force field^28,29^ in combination of TIP3P water,^30,31^ all converted to a format compatible with the GROMACS MD engine. ^32^

### Simulation methods

Here, we use a combination of parallel-tempering in the well-tempered ensemble and well-tempered metadynamics (PTWTE-WTM).^33–37^ For the (well-tempered) metadynamics component of this combined advanced sampling technique, we used a similarity-based order parameter to describe the similarity between a given configuration and a given reference state. The reference state is chosen as the native hybridized topology. In an ideal metadynamics sampling, the order parameter(s) used in order to accelarate the sampling should capture all slow degrees of freedom of given system. The order parameter that we use, the fraction of native-like contacts between strands, Q_*inter*_, accounts for inter-strand interactions such as base pairing, WC type hydrogen bonds, and cross-stacking interactions between strands. For other degrees of freedom that may not be directly captured by Q_*inter*_, such as intramolecular interactions, e.g. backbone torsional rotations, etc., we combine the (well-tempered) metadynamics with the parallel-tempering (in the well-tempered ensemble) scheme.

PTWTE^33,36^ part of PTWTE-WTM simulations are setup to cover a temperature range between 300 K and 475 K for the dodecamer and two of the decamers f(G·C)=0.7, 0.8); 279.5 K and 442.5 K for other decamers studied. In the framework of well-tempered ensemble, the potential energy is biased using 500 kJ/mol Gaussian width, 1.0 kJ/mol initial Gaussian height, with Gaussian potentials added every 2000 steps, with a bias factor of 16. Temperatures of the 14 replicas are distributed geometrically. The average replica exchange acceptance ratio is 25 % for decamers; 20% for the dodecamer.

WTM part of the simulations use only one order parameter, Q_*inter*_. Detailed description of the parameter is presented in the following section. The initial Gaussian height is set to 1.8 kJ/mol with a bias factor of 35 for the WTM sampling on Q_*inter*_. Gaussian width is set to 0.005 and since Q_*inter*_ is defined strictly between 0 and 1, lower and upper interval limits are applied,^38^ respectively at Q_*inter*_=0.01 and Q_*inter*_=0.99 together with lower and upper walls (harmonic potentials) with a spring constant of 75000 kJ/mol.

After initial solvation, all DNA systems are equilibrated in 100 ps NVT simulations (T=300 K) followed by 100 ps NPT simulations (T=300 K, P= 1 bar). Prior to starting the PTWTE-WTM simulations, unbiased NVT simulations of each replica are performed for 200 ps in order to equilibrate the potential energy of the replicas. Production simulations are run in NPT conditions where the temperature is maintained constant at given replica temperature using Nose-Hoover thermostat^39,40^ with a 1 ps time constant and at the atmospheric pressure (1 bar) using isotropic Parrinello-Rahman barostat^41,42^ with a time constant of 2 ps. Electrostatic interactions are calculated using the particle-mesh Ewald method^43^ with a real space cutoff distance of 1 nm. A cutoff distance is 1 nm is also used for the van der Waals interactions.

All simulations are performed using the GROMACS code (version 2016.3) ^44,45^ with the PLUMED (version 2.3.1) patch^46^ for metadynamics sampling.

### Description of the order parameter

Inspired by protein folding studies where folding reaction is described on the native-like contacts coordinate,^47–51^ we describe the DNA hybridization reaction on the inter-strand native-like atomic contacts. We define the order parameter Q_*inter*_ as 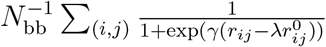 modifying the generalized definition of Q,^48^ for inter-strand contacts. The sum runs over N number of atomic pairs (i, j) that are considered in contact where the atom i is from strand A and atom j is from strand B (complementary strand). Any heavy (i.e. non-hydrogen) sidechain atom of strand A is considered in contact with that of strand B if the distance between them is less than 5Å in the reference native structure. 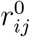 and *r*_*ij*_ are the distances between i and j in the reference native structure and in any given instantaneous configuration, respectively. *γ* in the smoothing function is taken as 0.5 nm and the fudging factor, *λ*, is taken as 1.5. Protein data bank (PDB) structure 1BNA is used as the reference native structure for DDD. Double-stranded B-DNA configurations of decamers are created by Nucleic Acid Builder (nab) tool of AmberTools ^52^ and used as the reference native structure in the description of Q_*inter*_.

### Two-state analysis

In this work, we assume hybridization reaction of DNA strands follows a two-state equilibrium:

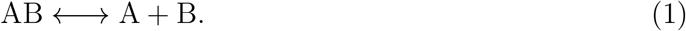

where the AB represents double-stranded helix (duplex); A and B are unhybridized, single strands, complementary to each other. We calculate the Δ*G*_*hyb*_ as the free energy difference between the hybridized state and the unhybridized states. Description of these states are explained in the Results and Discussion section.

The fraction of unhybridized states at the equilibrium is calculated as 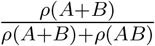 given that the probabilities, *ρ*, of states (A+B) and (AB) can be calculated from their relative free energies via Boltzmann relation. We define the melting temperature as the temperature at which *ρ*(*AB*) = *ρ*(*A* + *B*) = 0.5

Unbiased free energies (or potential of mean force [PMF]) are obtained after completion of biased simulations. Unbiased PMFs are recovered via the technique described by Tiwary and Parrinello. ^53^ Reweighting is performed considering all biases acting on potential energy and Q_*inter*_.

## Results and Discussion

We, first, characterize the free energy surface of Drew-Dickerson Dodecamer (DDD) DNA oligomer’s hybridization at 300 K. Sampling of Q_*inter*_ as a function of time/replica at 300 K can be seen in Figure S1. We identify an unhybridized basin where there are few native-like contacts, between Q_*inter*_=0 and Q_*inter*_=0.3 (Figure 1). After forming the initial native-like contacts, the free-energy goes downhill to as Q_*inter*_ increases, i.e. as more native-like contacts form, consistent with a nucleation-elongation mechanism as found by experiments and coarse-grained models of DNA.^11,12,54–56^

**Figure 1:**
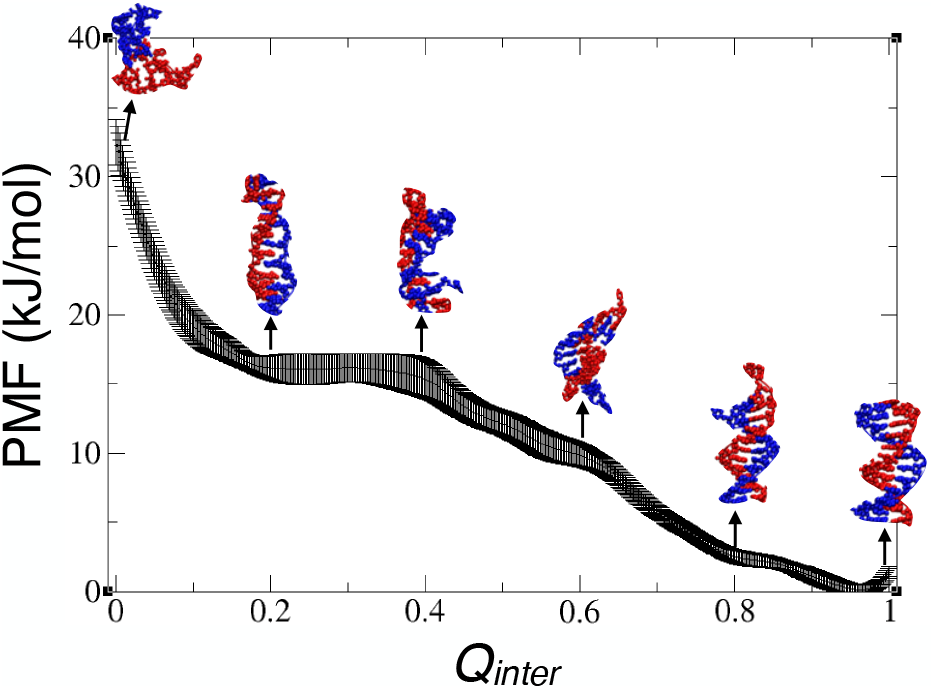
Free energy surface of hybridization of DDD oligomer at 300 K. Errors are calculated from the standard deviation of the last ten free energy profiles equally separated at every 10 ns. Snapshots are DDD oligomer (whose complementary strand has identical sequence) configurations at selected Q_*inter*_s as pointed by arrows.

### Thermal melting of Drew-Dickerson Dodecamer

We estimate the melting temperatures from PMFs (as function of Q_*inter*_) evaluated at various temperature (Figure 2A). We choose the unhybridized and hybridized states as the states Q_*inter*_=0.15 and Q_*inter*_=0.9, respectively. We then calculate the melting curve, i.e. fraction of hybridized DNA as a function of temperature from their relative free energies as described in the *Methods* in order to find the melting temperature. The melting curve is reported in Figure 2B after taking the finite-size effects into account^58^ and the melting temperature is found as 331.8 K.

**Figure 2:**
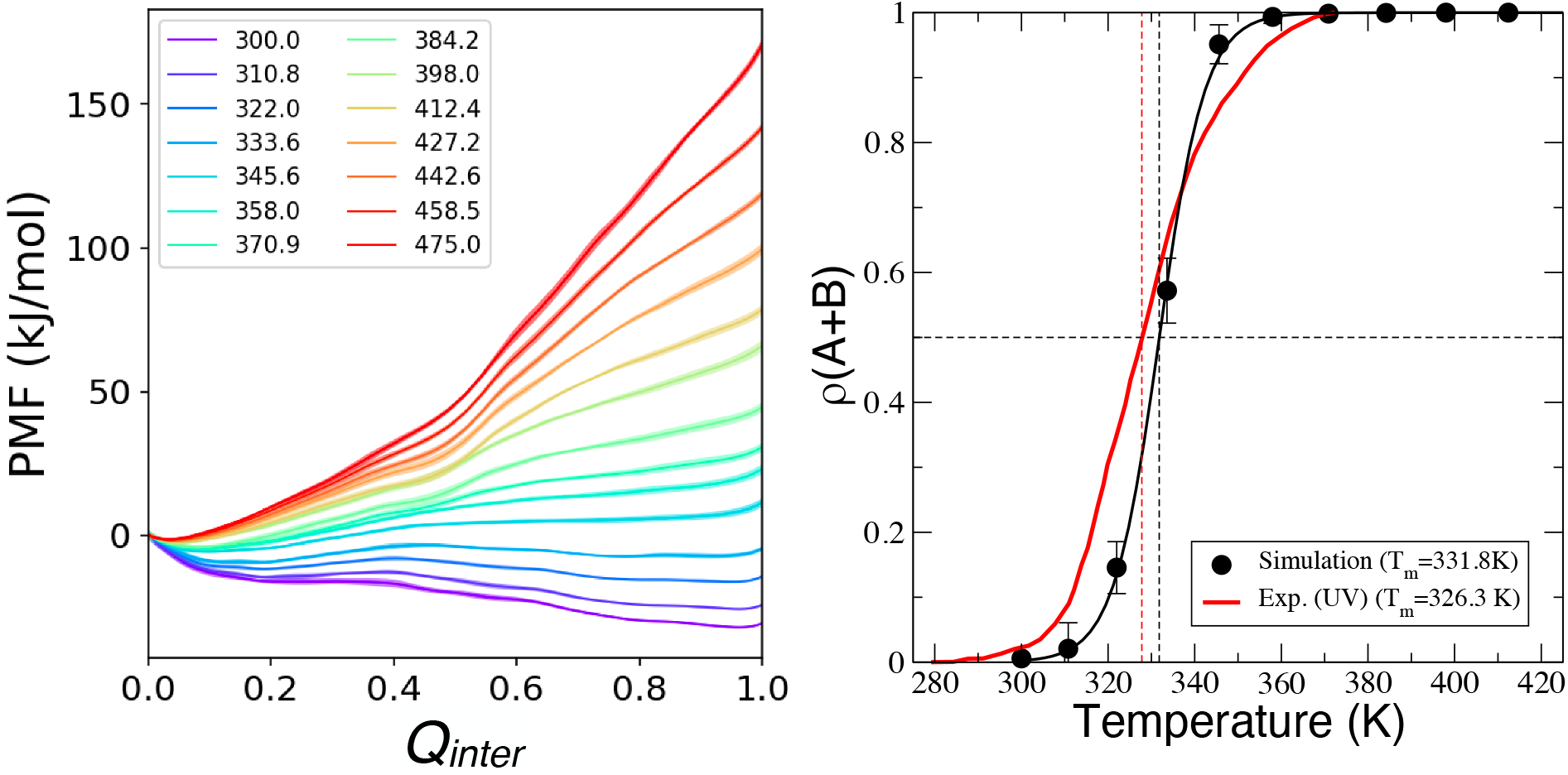
Thermal melting of DDD. Free energy surface of DDD oligomer is evaluated at several temperatures. Errors estimated from block averaging are shaded. (A.) Melting curve is calculated from respective free energies of hybridized and unhybridized states (B.) The temperatures at which *ρ*(*A* + *B*)=0.5 are designated as the melting temperatures. Experimental data is from thermal melting (UV) experiments.^57^

To ensure that we attain the equilibrium in sampling, we measure Δ*G*_*hyb*_ as a function of time (Figure S2) and consider the part of the trajectories where Δ*G*_*hyb*_ fluctuates within thermal fluctuation limits (±kT) for analysis. As a more stringent criterion of equilibrium, we also compare melting curves from two independent simulations (see Methods) where the initial conditions are unique, which show identical results for two simulations (Figure S3).

Comparison of our melting curve with the experimentally obtained one at the same salt concentration exhibits a good agreement between the melting experiment and our simulations (Figure 2B), with a caveat. Experimental data from Marky et al. ^57^ is collected at a strand concentration three orders of magnitude smaller than the DNA concentration in our simulation box. DNA concentration measurably affects the melting temperature^59^ as expected from the physical reaction of DNA hybridization being nonequimolar. As it is a non-negligible effect, strand concentration is further discussed in the next section where we systematically study the melting temperature of decamers.

The order parameter Q_*inter*_ is, not surprisingly, correlated with the number of native-like Watson-Crick (WC) type hydrogen bonds and number of stacked bases (Figure S4). To emphasize the importance of the order parameter selection for the success of sampling, we perform a supporting set of simulations using a similar combined advanced sampling. In this simulation, we have the same PTWTE-WTM setup, except for the order parameter. Instead of instead of Q_*inter*_, we use the number of native-like WC type hydrogen bonds as the order parameter. In Figure S5, we show that this sampling cannot accurately predict the melting. Indeed, during the entire course of the simulation, we never find that the hybridized state is more stable than the unhybridized state at 300 K (Figure S5). Therefore, we note that two order parameters being correlated does not necessarily mean that they perform equally successful in sampling. Q_*inter*_ outperforms the native-like WC type hydrogen bonds as Q_*inter*_ provides a more detailed description of hybrid DNA, which significantly reduces the degeneracy of states along the path between unhybridized and hybridized states.

We note that the unhybridized state, i.e. the state which only has small number of native-like contacts (low Q_*inter*_), not necessarily differentiates the configurations where two strands are far apart from each other and those where two strands are still relatively close to each other (although unhybridized). 2D reweighted free energy as a function of Q_*inter*_ and center of mass distance between two strands (Figure S6) shows that the configurations where two strands are considerably apart from each other are very high free energy configurations. Therefore, we emphasize that the hybridization free energy calculations in this work is related to the event of *melting* of native-like contacts, such as WC type hydrogen bonds, rather than pulling of strands apart from each other. Additional free energy calculations (umbrella sampling) that we perform show that pulling two strands fully apart from each other requires three to five times larger energy than just melting the native-like contacts at 300 K (Figure S7).

### Correlation between the experimental and simulation melting temperatures for varied GC content

To systematically show the generalized applicability of the method, we further perform simulations of DNA oligomers at various GC fractions, f(G·C), at fixed length (decamer) and fixed salt concentration (69 mM NaCl). We choose these oligomers as extensive experimental data is available for them. For the description of reference double-stranded helical (hybridized) state, we generate double-stranded helices of the given sequence in B-DNA configuration using nab tool of AmberTools^52^ (see *Methods*).

Following exactly the same methodology that we describe for DDD DNA in the previous section, we calculate the free energy surface of hybridization for various temperatures (Figure 3) and the melting temperatures for each oligomer. Comparison of melting temperatures calculated from our simulations and corresponding experimental melting temperatures are presented in Figure 4, for each f(G·C). Maximum difference between simulation and experiment fits in calculated to be less than ±15 degrees.

**Figure 3:**
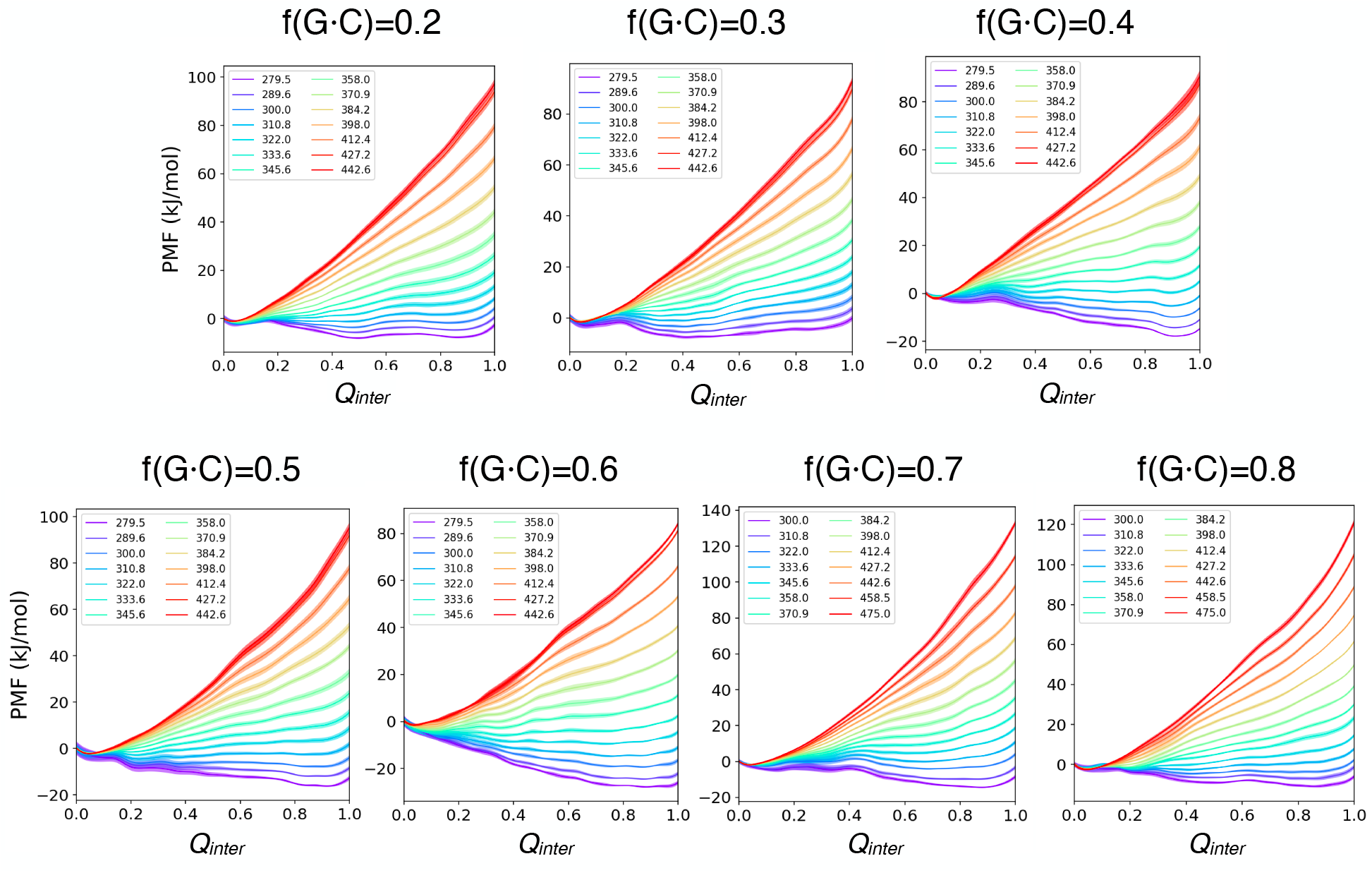
Free energy surface of DNA oligomers of various GC content, evaluated at several temperatures. Errors estimated from block averaging are shaded.

**Figure 4:**
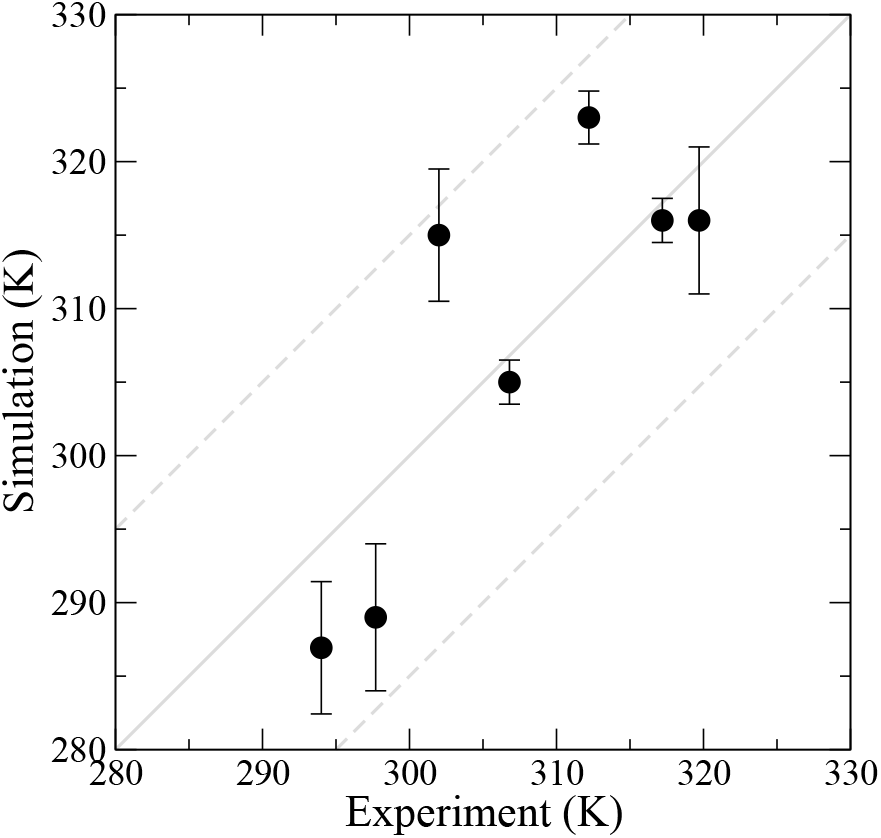
Correlation between the simulation and experimental estimates ^26^ of melting temperatures. Solid grey line is a guide to visualize perfect correlation (y=x) whereas the broken grey lines frame ±15°C deviation.

We note that the experimental melting temperatures are measured at 5 *μ*M strand concentration^26^ whereas our systems are several orders of magnitude more concentrated compared to the experiment (30 mM). An important downside of the practical application of this free energy calculation at atomistic resolution is the simulation cell size requirement. To keep the system computationally tractable at this scale, we are limited to concentrations on the order of mM; *μ*M concentrations in all atom, solvent explicit simulation are not possible to the current computational power. Melting temperature is expected to decrease upon strand dilution as there is a larger free volume available which can be occupied by unhybridized strands. Interestingly, however, we find the same free energy of hybridization and melting temperature upon dilution (twice larger volume) of f(G·C) = 0.7 DNA oligomer (melting curve illustrated in Figure S8).

To give a more quantitative idea on the effect of strand concentration on the melting temperature, we note that, for DDD DNA sequence in 100 mM NaCl solution, the melting temperature predicted from nearest neighbor model^5,6,60^ increases approximately by 5°C upon an order of magnitude increase in strand concentration. Given that the maximum difference between experimental estimates and our computational estimates from atomistic simulations could be as large as 15°C, it could be considered within error limits of our method, however, the strand concentration effect is still an area of future investigation.

### Comparison of hybridization thermodynamics for D-DNA and L-DNA chains

Lastly, we report that our method is symmetric, i.e. the ΔG_*hyb*_ calculated using our method for an L-DNA is the same as that for the D-counterpart of the DNA. We compare the melting curves of (D-)DDD DNA oligomer (naturally-occuring one) and (L-)DDD DNA oligomer (Figure 5) under the same salt and strand concentrations. We estimate approximately the same melting temperature for both chains. As the order parameters that we use in order to calculate FES relies solely on intermolecular distances (between complementary strands), it is not expected to be affected from the mirror symmetry. Accordingly, we use the same reference distances for (L-)DDD DNA as we use for (D-)DNA.

**Figure 5:**
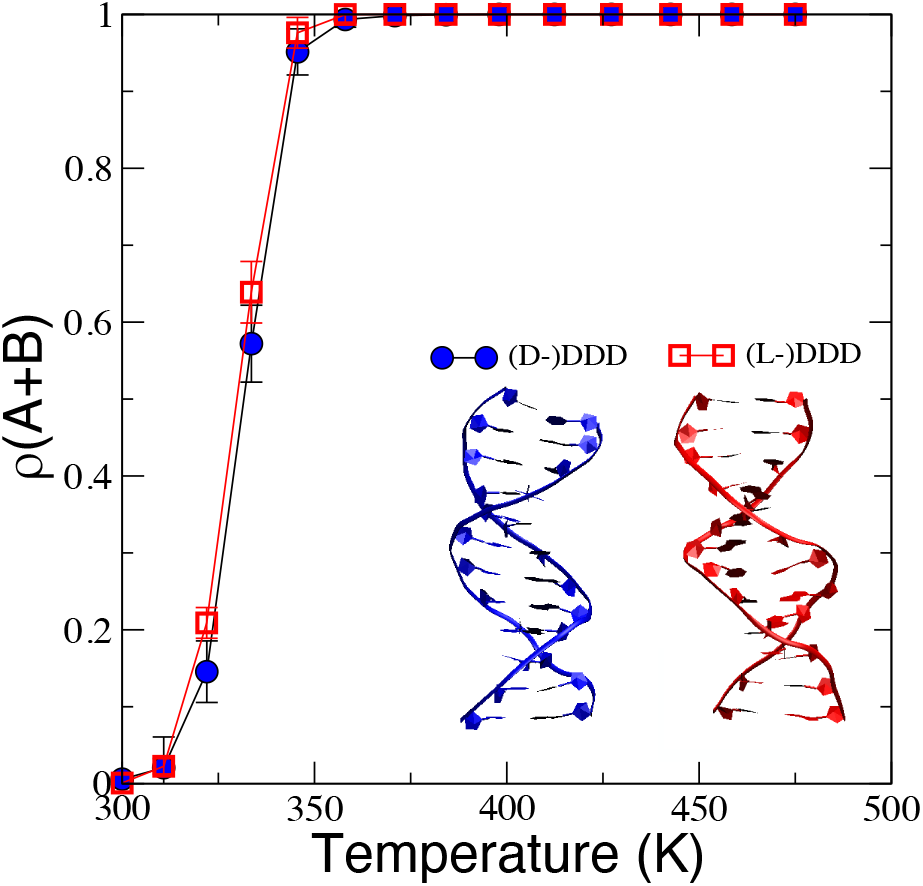
Comparison of melting curves for DDD at its two opposite chiral realizations. Blue structure is a representative double stranded helical structure of the hybridized state of (D-)DDD (right-handed helix) and the red structure is that of hybridized state of (L-)DDD, which is a left-handed helix (Q_*inter*_=0.99 for both).

Chirality of the DNA is imposed in the initial condition, which cannot change during the course of the simulation as the classical force fields do not allow bond breaking and forming. As there are no asymmetric terms in the force nucleic acid force field, we directly transfer all bonded and nonbonded interactions of D-nucleic acids to L-counterparts. As observed here, neither the free energy surfaces nor the melting curve is affected from the complete mirror symmetry, not surprisingly, as the underlying interaction parameters are exactly the same. The only difference is a total structural symmetry in the system, i.e. when hybridized, D-DNA forms of right-handed helix whereas the L-DNA forms a left-handed helix.

## Conclusions

For the first time, we quantify the hybridization free energy surface and melting profiles of short DNA oligomers at the atomistic resolution with a reasonable accuracy compared to the melting experiments. Using advanced sampling techniques and an appropriate order parameter, we push the limits of atomistic simulations, bypassing the need for a coarse-grained description of the hybridization event that takes place in milliseconds to seconds for given size of DNA oligomers.

Hybridization thermodynamics at the atomistic scale has been studied in more recent recent work, in addition to the work by Piana^22^ mentioned in the *Introduction*. In a study by Lomzov et al.,^61^ DNA melting temperature is calculated from the difference in the total energy of hybridized and that of unhybridized states, assuming that i) the volume change upon hybridization is negligible, ii) the difference in total energy of double-stranded state and that of single-stranded state is the same as the hybridization enthalpy when the assumption i) is true, and iii) hybridization entropy can be driven from a correlation between hybridization enthalpy and entropy. In another work, by Ren and coworkers, ^62^ relative hybridization free energy of DNA analogs have been estimated using an atomistic polarizable force field via alchemical transition method. However, we note that although the calculations in both work can report on melting temperature and thermodynamic quantities of hybridization, they cannot estimate the free energy surface of hybridization.

A particular power of our presented method arises naturally from it being a sampling technique based on metadynamics. Our method can be coupled with additional order parameters via, for example, concurrent metadynamics^37^ or parallel-biasing metadynamics.^63^ This coupling would be especially useful when there is a competing reaction of interest for a given nucleic acid system. An example of such a system may include a competition between surface adsorption of nucleic acid strands and hybridization. ^64^ Studying the equilibrium between these competing reactions would be useful to develop cancer sensing platforms in such a scenario as an example.

## Acknowledgement

GHZ thanks Daniel Kozuch and Dr. Pablo Piaggi for useful discussions. GHZ and PGD acknowledge the support from Unilever R&D. The simulations presented in this work are performed on computational resources managed and supported by Princeton Research Computing, a consortium of groups including the Princeton Institute for Computational Science and Engineering (PICSciE) and the Office of Information Technology’s High Performance Computing Center and Visualization Laboratory at Princeton University.

## Supporting Information Available

8 supporting figures are available.

This material is available free of charge via the Internet at http://pubs.acs.org/.

